# Spatial mutual nearest neighbors for spatial transcriptomics data

**DOI:** 10.1101/2024.10.08.615307

**Authors:** Haowen Zhou, Pratibha Panwar, Boyi Guo, Caleb Hallinan, Shila Ghazanfar, Stephanie C. Hicks

**Affiliations:** Bioinformatics and Systems Biology Graduate Program, University of California San Diego, La Jolla, CA, USA; School of Mathematics and Statistics, The University of Sydney, NSW 2006, Australia; Sydney Precision Data Science Centre, University of Sydney, NSW 2006, Australia; Charles Perkins Centre, The University of Sydney, NSW 2006, Australia; Department of Biostatistics, Johns Hopkins Bloomberg School of Public Health, MD, USA; Department of Biomedical Engineering, Johns Hopkins University, Baltimore, MD, USA; Center for Computational Biology, Johns Hopkins University, Baltimore, MD, USA; Malone Center for Engineering in Healthcare, Johns Hopkins University, MD, USA

**Author notes:** Co-corresponding authors: Shila Ghazanfar and Stephanie Hicks.

**Keywords:** spatial transcriptomics, unsupervised clustering, mutual nearest neighbors

## Abstract

Mutual nearest neighbors (MNN) is a widely used computational tool to perform batch correction for single-cell RNA-sequencing data. However, in applications such as spatial transcriptomics, it fails to take into account the 2D spatial information. Here, we present *spatialMNN*, an algorithm that integrates multiple spatial transcriptomic samples and identifies spatial domains. Our approach begins by building a *k*-Nearest Neighbors (kNN) graph based on the spatial coordinates, prunes noisy edges, and identifies niches to act as anchor points for each sample. Next, we construct a MNN graph across the samples to identify similar niches. Finally, the spatialMNN graph can be partitioned using existing algorithms, such as the Louvain algorithm to predict spatial domains across the tissue samples. We demonstrate the performance of spatialMNN using large datasets, including one with *N*=36 10x Genomics Visium samples. We also evaluate the computing performance of spatialMNN to other popular spatial clustering methods. Our software package is available at (https://github.com/Pixel-Dream/spatialMNN).

## 1 Introduction

Recent advances in spatially-resolved transcriptomics (SRT) have transformed the landscape for how we measure gene expression in intact tissue [1, 2]. These technologies can profile gene expression at a sub-, near-, or cellular resolution depending on the spatial technology used [3, 4]. Unsupervised clustering has become a standard data analytic step in the analysis of SRT data where the aim is to partition spatial units, cells for image-based technologies or spots for sequencing-based technologies, measured in the 2-dimensional space into either cell types or spatial domains. These cell types or spatial domains can be further explored in downstream analyses, like using differential expression across spatial domains to identity key marker genes. As SRT technologies are used to profile tens to hundreds tissue sections from different samples, an important challenge is to integrate multiple SRT samples across technical batches and simultaneously identify consistent spatial domains in population- or atlas-scale datasets [5].

Currently, there are two categories of existing computational methods to integrate multiple SRT samples: (i) methods that were developed to integrate multiple single-cell RNA-sequencing (scRNA-seq) samples across batches [6–10], and (ii) more modern methods that were developed for the analysis of SRT data by taking into account additional features beyond gene expression counts, including spatial coordinates or image features [11–17]. While the first category of robust methods are widely used in practice for the analysis of scRNA-seq data, they fail to leverage the spatial information, which has shown to be important for downstream analyses [18–20]. In addition, these more modern tools can be technology-dependent and computationally inefficient, including both running time and memory-usage [19, 20]. Finally, existing benchmark evaluations to spatially cluster SRT samples have primarily evaluated performance within a tissue section or utilize a method such as PASTE [21] to physically align two tissue sections before performing spatial clustering.

To address these problems, we developed spatialMNN, an algorithm that integrates multiple SRT samples using graph-based approaches and then identifies spatial domains across the SRT samples jointly. Our approach leverages mutual nearest neighbors (MNN) [8], which is used by many computational tools [7, 22, 23] and has consistently been recognized for its high performance in both accuracy and scalability in terms of memory efficiency and speed [10]. We demonstrate the performance of spatialMNN using both simulated and real SRT data sets, which scales to large-scale SRT datasets. We also evaluate the computing performance of spatialMNN to other popular spatial clustering methods.

## 2 Results

### 2.1 spatialMNN for batch correction and detecting spatial domains

In a similar spirit to MNN [8], the spatialMNN algorithm is designed to integrate multiple transcriptomics datasets, except spatialMNN is designed for transcriptomics datasets with spatial coordinate information, which can be used in downstream analyses to identify shared spatial domains (or clusters) across the samples (**Figure 1, Extended Data Figure 1**). Broadly, our approach begins by building a kNN graph based on the spatial coordinates within each tissue sample. Specifically, we denote the kNN graph as *G* =*< V, E >*, where *V* represents the set of all spots/cells and *E* the set of edges connecting each node *u* ∈ *V* with its *k* nearest neighbors. Using edge weights measured by Pearson correlation or the number of Shared Nearest Neighbors (SNN) on gene expression, we prune noisy edges using a novel smoothed edge pruning algorithm (**Extended Data Figures 2 and 3**, described in greater detail in **Section 4.1**). Using these pruned edges, spatialMNN identifies niches to act as *anchor points* for each sample. Next, we construct a MNN graph across the samples to identify similar niches across the samples. Finally, the spatialMNN graph can be partitioned using existing algorithms, such as the Louvain algorithm, to predict spatial domains across the tissue samples.

**Figure 1:**
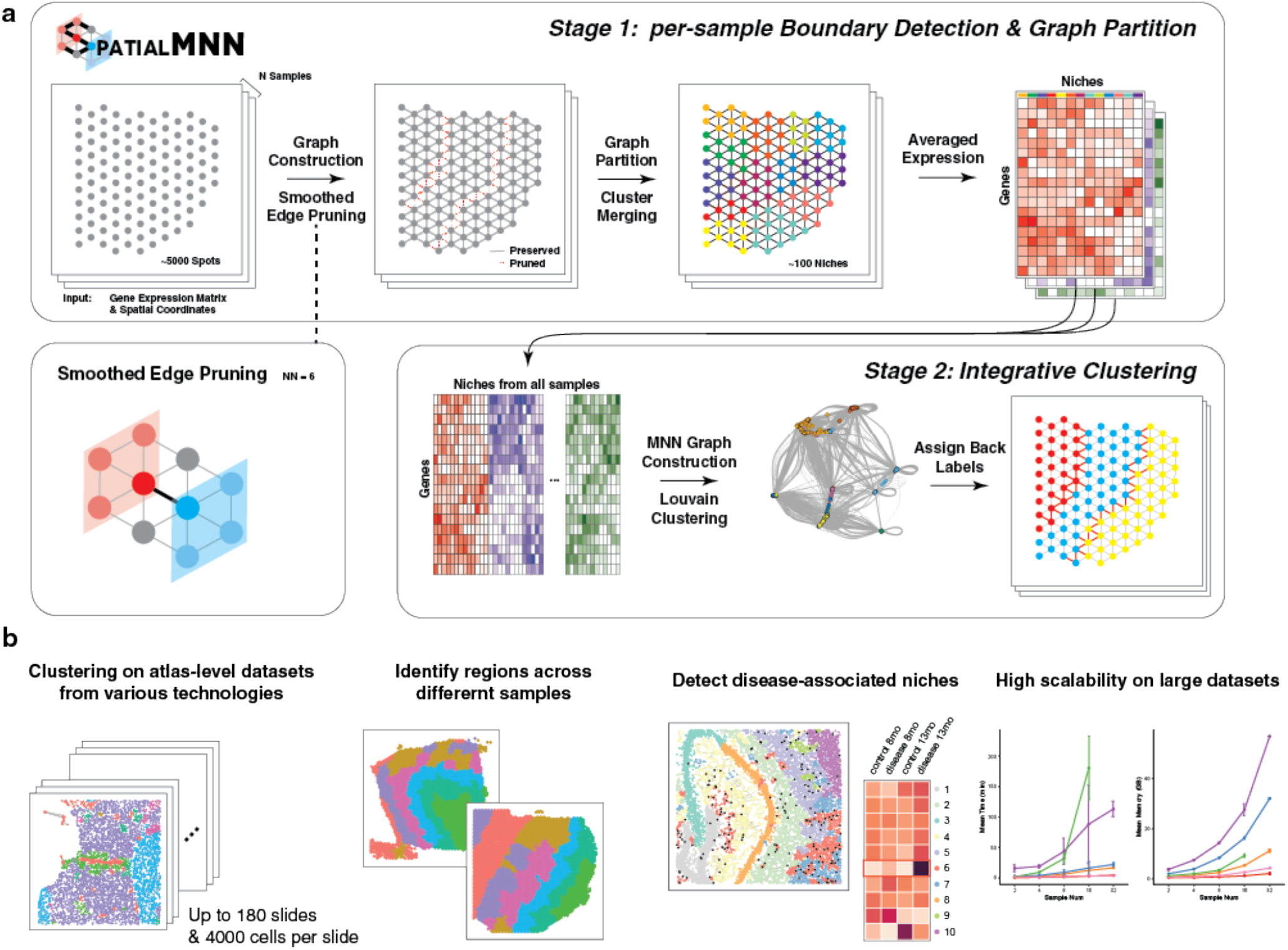
Overview of spatialMNN to integrate multiple SRT samples and perform downstream analyses including spatial domain detection. **(a)** Given a set of *N* multi-sample SRT datasets, the spatialMNN algorithm builds a *k*-Nearest Neighbor (kNN) graph based on the spatial coordinates and gene expression within each tissue sample. Next, edge weights are smoothed (considering neighboring spots/cells) and pruned to identify a set of *anchor points* for each sample. Then, spatialMNN constructs a MNN graph across the samples, followed by Louvain clustering to identify similar niches across the samples using gene expression that has been averaged across spots/cells within a niche. The resulting clusters are assigned back to the original spatial coordinates. **(b)** spatialMNN can be used in downstream analyses, including detecting spatial domains in large-scale atlas datasets, identifying regions across different samples, and detecting disease-associated niches. We demonstrate how spatialMNN is highly scalable and accurate on large datasets.

### 2.2 Key innovations of spatialMNN

The key innovations of spatialMNN compared to existing approaches are as follows. First, recent work [5] described challenges to integrate multiple SRT samples, including how existing approaches that perform multi-sample batch correction for SRT use existing methods developed for scRNA-seq data [7–10]. A key innovation of spatialMNN is that it takes into account spatial coordinates as input, in addition to the gene expression, making it uniquely designed for SRT data. Second, due to spatialMNN being a graph-based approach, it can be applied to either non-targeted RNA capture and sequencing, such as Slide-seq or 10x Genomics Visium, or image-based, targeted, *in situ* transcriptomic profiling at a molecular and single-cell resolution, such as MERFISH or Xenium [4]. Third, because a divide-and-conquer approach of first identifying spatial niches within each sample separately is used, the computational complexity of spatialMNN is reduced and runtime is fast. Specifically, the partitioning of the graph into anchor points (or niches) serves as a form of data reduction, where the information from potentially thousands of spots/cells is reduced to hundreds of anchor points. Similarly, SpaceFlow-DC [19, 24] also uses a divide and conquer approach, while other methods are designed for one sample at a time or are infeasible to apply to large datasets. As the cost of generating these data decrease, computational complexity and runtime to integrate SRT data across multiple tissues or individuals will become increasingly important to perform population-level analyses [5].

### 2.3 spatialMNN outperforms existing methods in both accuracy and scalability

To demonstrate the applicability of spatialMNN on real SRT datasets, we considered three datasets across different technologies and scales. In the first dataset, Wang et al. [25] used the STARmap platform to profile the prelimbic area in mouse brain across 3 tissue sections, 166 genes, and 3,190 cells (**Figure 2a-d, Extended Data Figure 4**). The second dataset from Maynard et al. [26] used the 10x Genomics Visium Spatial Gene expression platform to profile postmortem human dorsolateral prefrontal cortex (DLPFC) across 12 tissue sections, 33,538 genes, and 47,681 spots (**Figure 2e-h, Extended Data Figure 5**). In the third dataset, Joung et al. [27] used the MERFISH platform to profile the mouse frontal cortex and striatum across 31 tissue sections, 374 genes, and 378,918 cells (**Figure 2i-l, Extended Data Figure 6**). In each of these datasets, the manually labeled cell type or spatial domains (which we use as approximate ground truth) are used to help compare the performance of spatialMNN to existing algorithms (labeled ‘Ground Truth’ in **Figure 2a,e,i**). We applied spatialMNN to integrate the multiple tissue sections within each dataset and compared to existing algorithms that can be used to detect spatial domains across multiple samples, including BASS [13], PRECAST [14], BayesSpace [11], BANKSY [15], SLAT [16], MENDER [17] and Louvain [6]. We used the adjusted Rand index (ARI) to evaluate the spatial domains detected by each of the algorithms compared to the manually labeled cell types or spatial domains. We further evaluated the runtime and memory used over the course of the algorithm.

**Figure 2:**
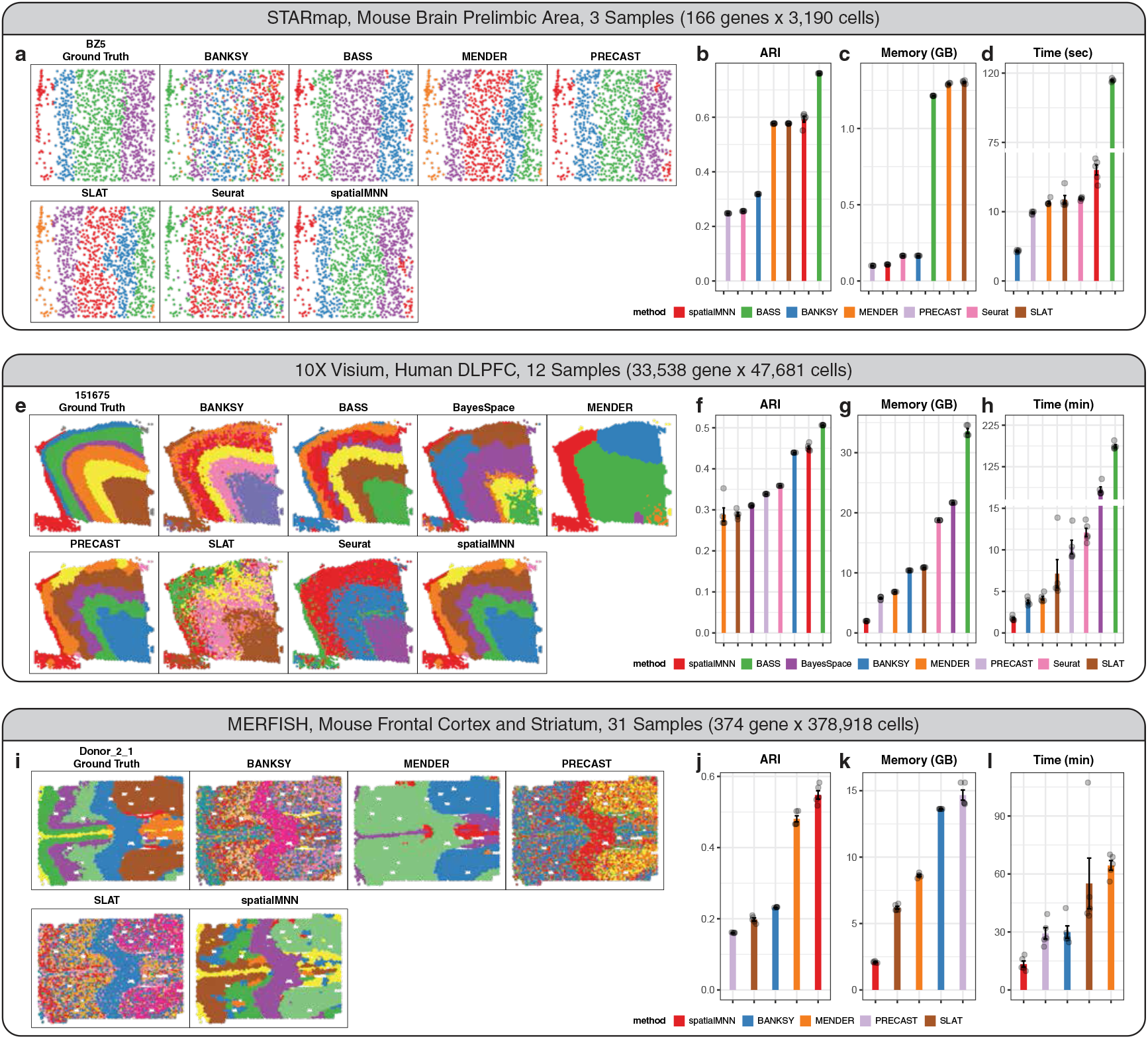
Application of spatialMNN to integrate multiple tissue sections and detect cell types and spatial domains across samples within three datasets measured on the STARmap, Visium, and MERFISH platforms. We considered three datasets (**a-d**) Wang et al. [25] used STARmap with *n*=3,190 cells across *N*=3 tissue sections measured on the STARmap platform, (**e-h**) Maynard et al. [26] with *n*=47,681 spots across *N*=12 tissue sections measured on the 10x Genomics Visium platform, and (**i-l**) Joung et al. [27] with *n*=378,918 cells across *N*=31 tissue sections measured on the MERFISH platform. **(a, e, i)** Visualization of one tissue section from each dataset colored by the manually labeled (‘Ground Truth’) or predicted cell types/domains from spatialMNN, BASS [13], PRECAST [14], BayesSpace [11], BANKSY [15], SLAT [16], MENDER [17] and Louvain [6]. Performance evaluation (*y*-axis) of **(b, f, j)** adjusted Rand index (ARI) comparing the performance of the algorithms (*x*-axis) to the manual annotations (repeated 5 times), **(c, g, k)** elapsed time (minutes) for each of the algorithms (*x*-axis), **(d, h, l)** maximum memory (RAM) used (GB) for each of the algorithms (*x*-axis). Louvain and BASS are excluded in MERFISH frontal cortex benchmark due to out-of-memory issue. BayesSpace is excluded in STARmap brain prelimbic area and MERFISH frontal cortex benchmarks due to application limitation. All benchmarks are run on a high performance computing cluster (8 cores, 128GB max memory and 36 hrs max running time).

While BASS outperformed spatialMNN in accuracy compared to the manual labels (assessed with ARI) in the first and second dataset (**Figure 2b,f,j**), we found the runtime and memory usage requirements for BASS to be significantly higher compared to existing algorithms (**Figure 2c,d,g,h**). Specifically, in the first dataset with 3,190 cells across 3 tissue sections, BASS took 115 seconds to complete comparedto spatialMNN, which took 16 seconds. Similarly, in this dataset, BASS required 1,243 MB of memory compared to 110 MB with spatialMNN. In the second dataset with 47,681 spots across 12 tissue sections, BASS and spatialMNN used 175 and 1.7 minutes, along with 33.6GB and 1.98GB, respectively. In the third dataset, we were unable to run BASS entirely due to out-of-memory issue (maximum 128GB, **Figure 2k,l**) while on average, spatialMNN only used about 2GB of memory and 15 minutes to finish the clustering. Finally, when comparing spatialMNN to other existing algorithms, we found that spatialMNN resulted in the highest ARI score along with a quicker running time and smaller memory-usage.

### 2.4 spatialMNN identifies disease-associated niches in Alzheimer’s Disease

Understanding the specific patterns of the spatial organization of cells can be important to understand the development and the pathology of disease, such as cancer or Alzheimer’s Disease (AD). However, existing algorithms for mining these spatial patterns rely on transcriptome-based cell type annotation [29, 30]. Here, we used spatialMNN in an unsupervised manner to identify disease-associated niches in mouse Alzheimer’s Disease (AD) samples [28].

Using spatialMNN, we identified unique spatial domains and cell types within mouse brain regions across control and AD tissue (**Figure 3a,b**). By comparing the clustering results of AD samples from 13-month (a) and 8-month (b) mice, we observed that cluster 6 emerged as a disease-associated niche, showing a unique spatial distribution pattern related to AD pathology. A heatmap comparing cluster abundance across different samples highlights the increase of cluster 6 in AD samples, especially in the 13-month sample (**Figure 3c**). Specifically, we found a progressive enrichment of this disease-associated niche with increasing age and disease severity.

**Figure 3:**
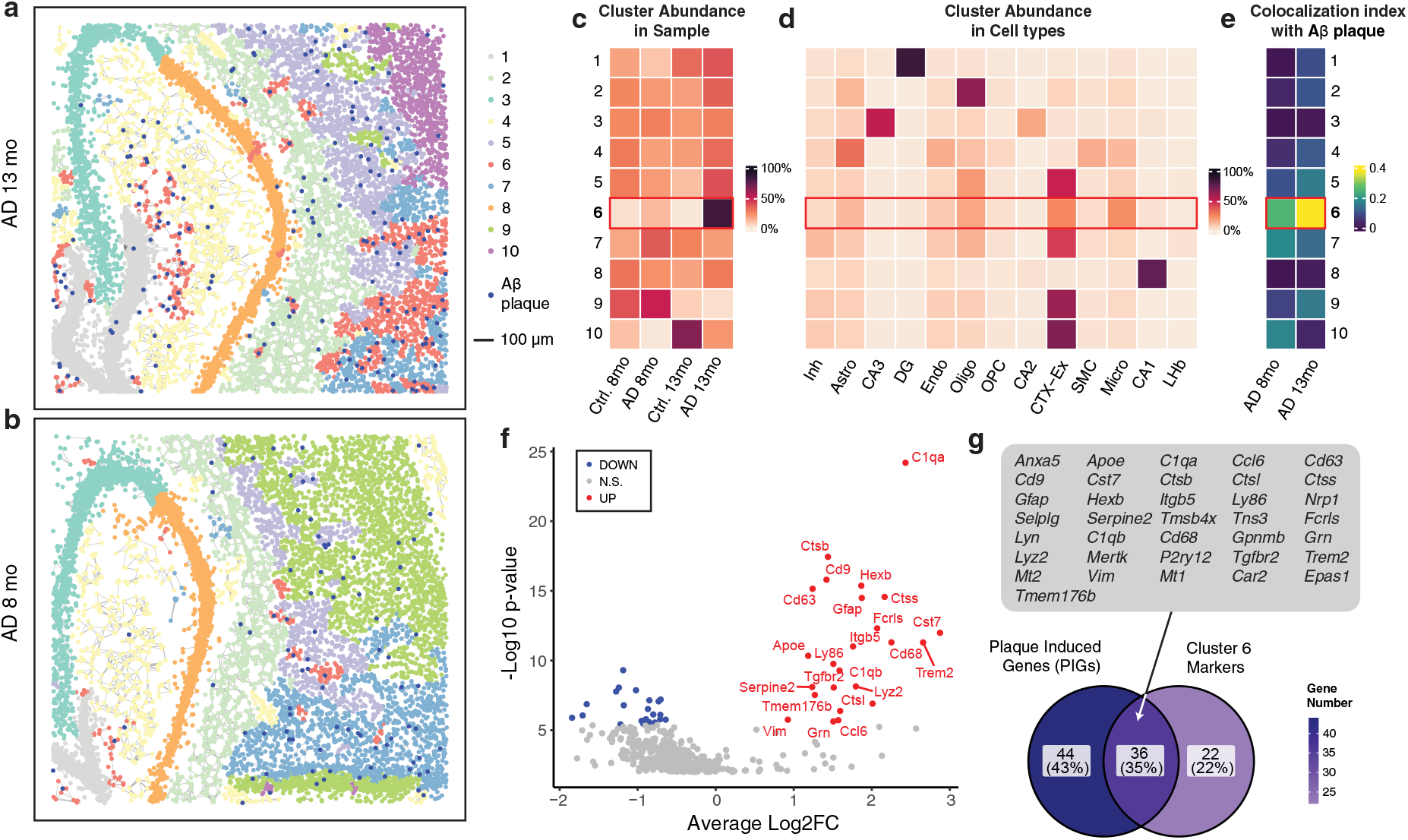
SpatialMNN identifies disease-related niches in mouse Alzheimer’s Disease samples. STARmap PLUS was used to profile the gene expression and protein (in the same tissue) from *N*=4 mouse samples (Control (*N*=2) and Alzheimer’s Disease (AD) (*N*=2) mice at ages 8 months (mo) and 13 mo) [28]. **(a, b)** To integrate and perform multi-sample spatial clustering, spatialMNN identified 10 predicted niches (or unsupervised clusters) in two AD samples at **(a)** 13 months and **(b)** 8 months, with dark blue spots indicating A*β* plaques (protein). The plotting scale is shown in the legend. **(c)** Heatmap of the cluster abundance across samples, highlighting the increased presence of cluster 6 in AD samples (compared to control samples), especially in the 13 mo sample, suggesting cluster 6 is a disease-associated niche. **(d)** Heatmap of the cluster abundance across cell types, suggesting the microglia (Micro), astrocytes (Astro), and oligodendrocytes (Oligo) cell types are enriched in cluster 6. **(e)** Co-localization index heatmap shows the spatial correlation of clusters with A*β* plaques, with cluster 6 having the highest co-localization index. **(f)** Differential expression (DE) analysis identifies marker genes in cluster 6. **(g)** Venn diagram comparing the marker genes found in cluster 6 with the Plaque Induced Genes (PIGs) identified in the original study, showing a significant overlap. All overlap genes were annotated in the gray box.

Using published cell type annotations in the original paper [28], we found differences in the cell type abundances within the spatialMNN clusters (**Figure 3d**). Specifically, we found that cluster 6 is enriched for microglia, astrocytes, and oligodendrocytes. This suggests that these cell types are key components of the identified disease-associated niche, aligning with literature findings [31] and underscoring the qualitative accuracy of spatialMNN in identifying disease-associated niches. Next, we investigated if the predicted clusters were co-localized with the location of the A*β* plaques using a spatial correlation metric. The spatial co-localization analysis found that cluster 6 resulted in the highest co-localization index with the A*β* plaques, further emphasizing the relevance of cluster 6 to AD pathology (**Figure 3e**).

Finally, we used differential expression (DE) analysis to identify marker genes for cluster 6 (**Figure 3f**). We compared top DE genes from this cluster with Plaque Induced Genes (PIGs), identified in the original study, and found strong overlap between these gene sets. This suggests consistency between the identified marker genes and previously reported PIGs (**Figure 3g**). Overall, we found spatialMNN effectively identified disease-associated niches in the spatial transcriptomics dataset, revealing key spatial and molecular aspects of Alzheimer’s Disease pathology in mouse models.

### 2.5 spatialMNN in application to atlas-scale datasets

To demonstrate how spatialMNN can scale to datasets with a larger sample size, we used a dataset with *N*=31 tissue sections from Nelson et al. [32] who used the 10x Genomics Visium Spatial Gene Expression platform to profile postmortem human hippocampus across ten donors, 31,483 genes, and 150,917 spots. Using spatialMNN, we integrated and identified 13 spatial domains across the *N*=32 tissue sections. We found the spatialMNN spatial domains were consistent with domains found by the original authors(**Figure 4a,b**). Specifically, in the original paper, the authors manually annotated the spots based on known morphology and gene markers along with applying PRECAST [14] to annotated spots using a data-driven approach to a set of “broad domains.”

**Figure 4:**
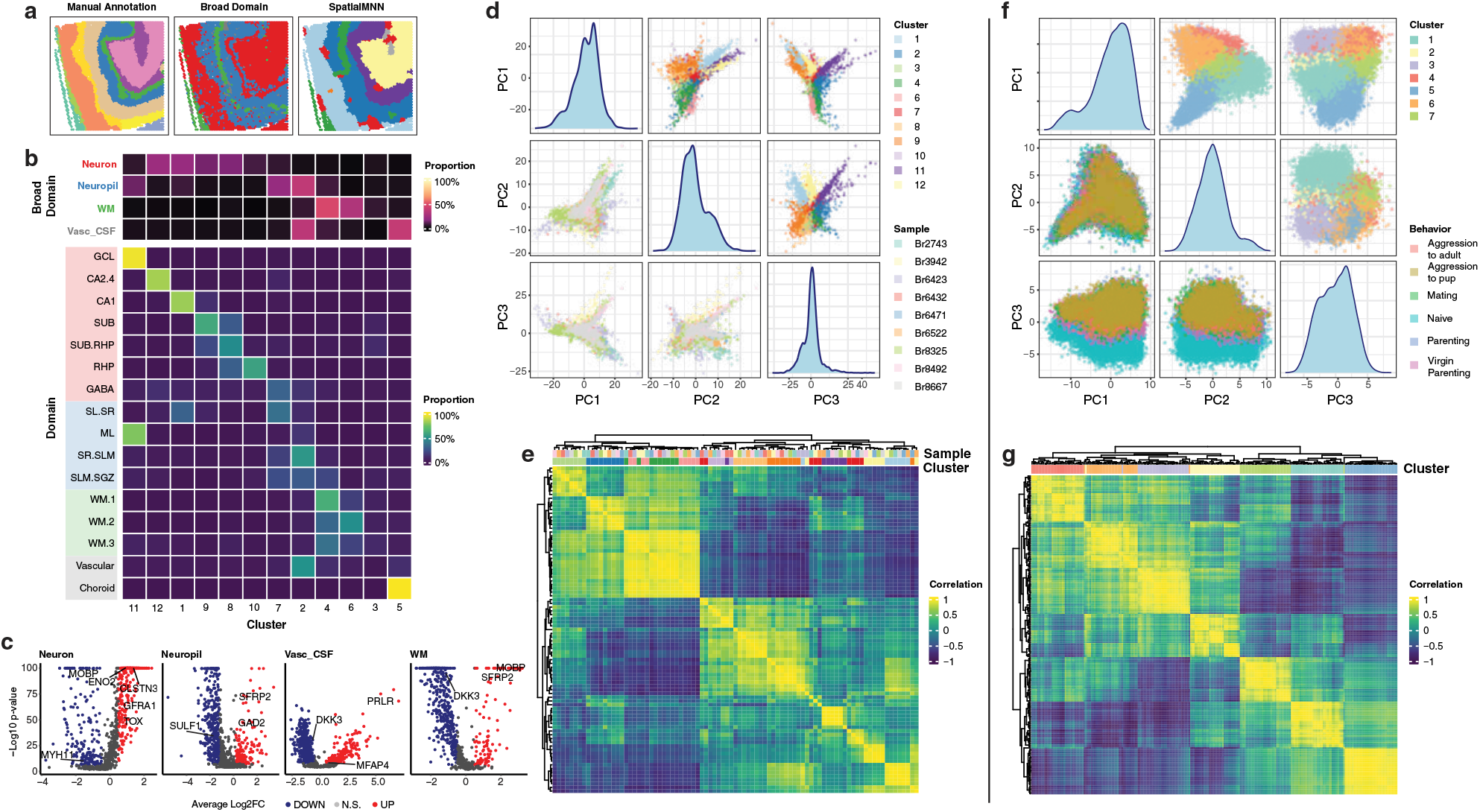
Application of spatialMNN to large SRT datasets with *N*=32 Visium samples and *N*=181 MERFISH samples. Panels (a-e) and (f-g) are based on *N*=31 tissue sections from human hippocampus profiled on the 10x Visium platform [32] and *N*=181 slides (over 870,000 cells) profiled on MERFISH [33], respectively. **(a)** Spot plots of sample *Br8667_V11L05-336_A1* ‘s domains using a manual annotation, a data-driven approach to “broad domain”, and spatialMNN. **(b)** Heatmap showing the fraction of different cell types/broad domain types in each cluster identified by spatialMNN **(c)** Volcano plots of four broad domains, highlighting differentially expressed genes across identified clusters overlap with the manually annotated domains: Neuron (clusters 1, 8, 9, 12), Neuropil (clusters 4, 8, 12), White Matter (WM; clusters 3, 4, 6), and Vascular CSF (cluster 2, 5). **(d)** Distribution of identified clusters and samples in the top three principal components (PCs). **(e)** Heatmap of Pearson correlation demonstrating the consistency of identified clusters based on gene expression profiles. **(f)** Distribution of identified clusters and sample-level variables projected along the top three PCs, similar to panel (d), showing the robustness of spatialMNN in a larger dataset. **(g)** Correlation heatmap of identified clusters.

Motivated by the use of the defined broad domains by Nelson et al. [32], we similarly combined the spatial domains detected by spatialMNN to a comparable resolution of “neuron” (clusters 1, 8, 9, 12), “neuropil” (clusters 4, 8, 12), “white matter” (WM; clusters 3, 4, 6), and “vascular CSF” (cluster 2, 5). We identified DE genes within these broad domains from spatialMNN and found a consistent set of DE genes from the original authors (**Figure 4c**). Using the predicted domains from spatialMNN, we performed principal components analysis (PCA) and found that the spatialMNN domains explained the most variation in the top three principal components (PCs), compared to the donor-level variation (**Figure 4d**). Furthermore, we found strong correlation at the gene-expression level across the samples within each cluster (**Figure 4e**).

In a second example, we considered a dataset consisting of *N*=181 slides with over 870,000 cells profiled on the MERFISH platform [33]. Similar to the first dataset, we used spatialMNN to detect 7 spatial domains jointly across all 181 slides. We found the spatialMNN domains explained the most amount of variation compared to other known variables (**Figure 4f**). Finally, we found strong correlation at the gene-expression level across the cells from different samples within each cluster (**Figure 4g**).

### 2.6 spatialMNN is fast and memory-efficient

In addition to evaluating spatialMNN using real SRT samples, we also simulated SRT samples to demonstrate how spatialMNN scales with both running time and memory efficiency. We considered SRT datasets ranging from *N*=2-64 samples, each with a known set of spatial domains made up as a mixture of cell types within each spatial domain (**Figure 5a**). Briefly, we developed a two-stage approach for simulating SRT data (see **Section 4.2.7** for more details), starting with four predefined spatial patterns (**Figure 5b**) designed to mimic different types of tissue spatial distributions. Next, we simulated the number of cells in each spot using a uniform distribution. Then, we assigned cell types to each cell based on a pre-defined tissue-to-cell type probability table (**Figure 5c**). Each cell type was assigned 200 marker genes, resulting in a total of 800 marker genes. To simulate realistic conditions, we also incorporated 10% noisy genes. Finally, for each simulated sample, the resulting gene expression matrix consisted of 889 genes and 4,992 cells simulated from Gaussian distributions. We generated a series of seven datasets, with sample sizes of *N*=2, 4, 8, 16, 32, 48, and 64 SRT samples, and the number of spots ranging from approximately 10,000 to 320,000. In this way, the simulated data is representative of current computational challenges posed by existing large-scale datasets.

**Figure 5:**
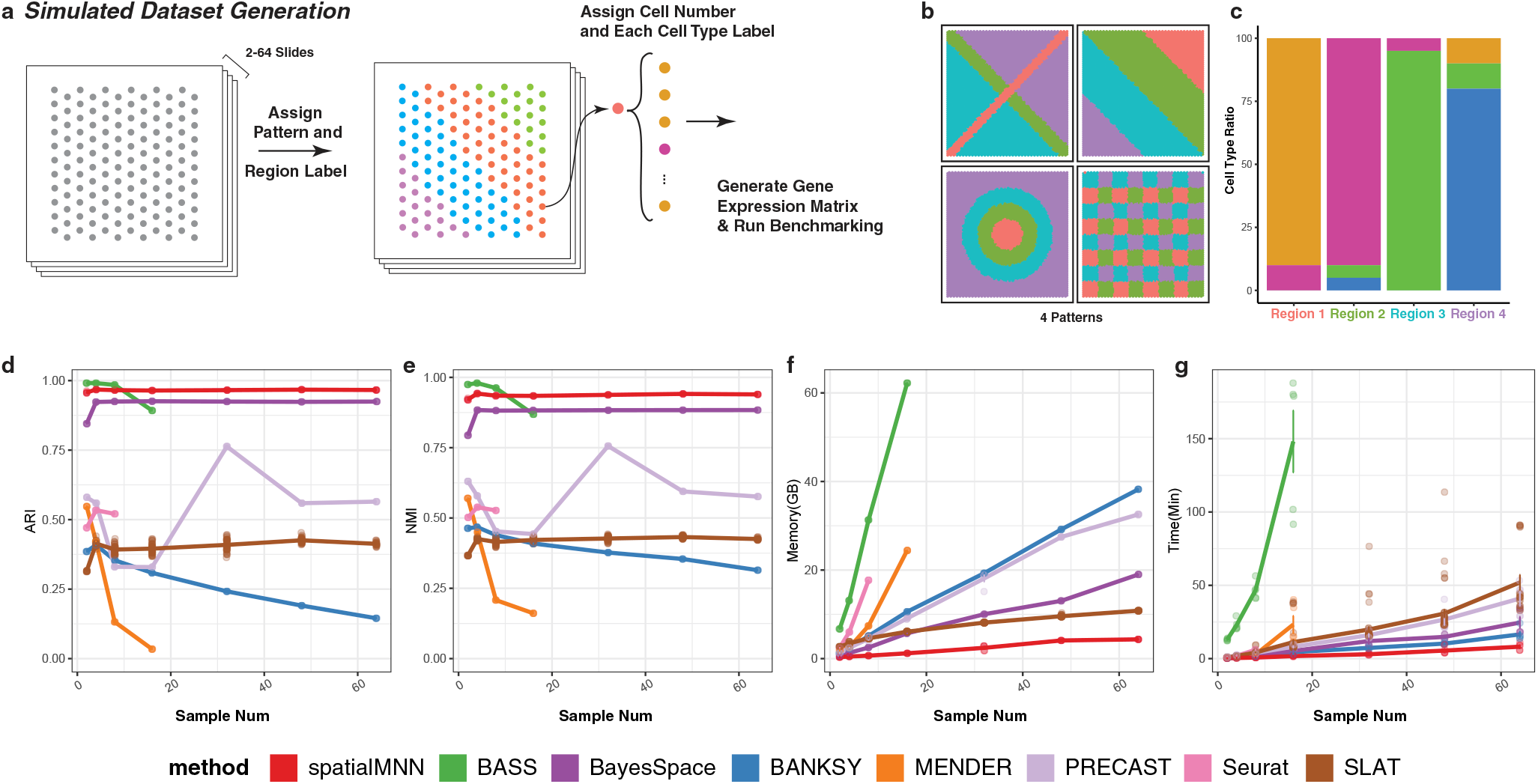
Simulation benchmark reveals the low memory and time efficiency of SpatialMNN. **(a)** Overview of the simulated dataset generation process: patterns and region labels are assigned to cells across 2-64 slides, with cell numbers and types specified to generate a gene expression matrix for benchmarking. **(b)** Illustration of the four distinct spatial patterns used in the simulation. **(c)** Bar plot showing the cell type composition across four different regions. **(d-g)** Benchmarking results comparing spatialMNN with other methods (BASS, Banksy, BayesSpace, MENDER, PRECAST, Seurat, and SLAT) on **(d)** Adjusted Rand Index (ARI), **(e)** Normalized Mutual Information (NMI), **(f)** memory usage, and **(g)** runtime across different sample sizes. Each simulated benchmark with different sample size was repeated 5 times. SpatialMNN demonstrates high clustering accuracy with consistently high ARI and NMI scores, while maintaining significantly lower computational costs in terms of runtime and memory usage.

Using our simulated SRT data, we found that spatialMNN is both accurate, based on ARI and normalized mutual information (NMI), (**Figure 5d-e**) and memory efficient (**Figure 5f**), as spatialMNN can identify spatial domains from *N*=64 SRT samples with less than 5GB of memory (RAM). Furthermore, spatialMNN is computationally fast as it can identify spatial domains from *N*=64 SRT samples in less than 10 minutes. Compared to existing methods, we found spatialMNN outperforms other approaches in all three categories, including accuracy, memory usage, and running time. Notably, while the accuracy performance of other methods like BASS, BANKSY and MENDER declines as the number of samples increases, spatialMNN maintains robust accuracy. In terms of computational efficiency, spatialMNN significantly outperforms other methods, exhibiting lower memory usage and shorter runtime (**Figure 5f,g**). Methods like BASS, Seurat and MENDER display a pronounced increase in either memory consumption or computational time, and sometimes both with larger datasets, whereas spatialMNN scales more efficiently. These results highlight the ability of spatialMNN to deliver high clustering accuracy with minimal computational cost, making it an effective and scalable solution for large-scale spatial transcriptomics analysis.

## 3 Discussion

Here we introduce spatialMNN, a new approach to integrate multiple SRT data and identify spatial domains across all samples. The computing performance of spatialMNN is greatly increased compared to other algorithms due to its reduced computational complexity via divide-and-conquer strategy, as well as the straightforward implementation of parallelization in the first stage of spatialMNN. To reduce the memory burden even further, future work could focus on the use of delayed operations to utilise data stored on-disk rather than in memory. As the accessibility to SRT technologies increase, the scale of datasets will inevitably rise too. Regardless, we have shown that spatialMNN can scale well for these datasets, as shown by the method’s strong performance on one of the largest publicly available SRT datasets

The spatialMNN approach results in spatial domains that are consistent across multiple samples. Therefore, no per-sample clustering to *post-hoc* matching of clusters across samples is needed, ultimately reducing the burden of manual examination by analysis. Furthermore, the abundance of spatial domains identified can be used to extract a deeper understanding of a set of samples, as we showed with the AD dataset (**Figure 3**), which could be used to identify disease-related niches. In particular, when spatialMNN is applied to point-based SRT datasets, the clusters identified are actually spatial domain types. These are regions where a certain distribution of gene expression is observed, which can arise due to distinct demographics of cell type mixtures. Hence, it is important to note that the spatial domains found in spatialMNN will not always correspond to cell type identities.

Overall, spatialMNN is a novel approach that utilizes mutual nearest neighbors and was inspired by the need for less time-intensive and more accurate clustering methods for multi-sample SRT datasets. With increase in the number of large-scale SRT datasets, it will be important to ensure we can perform unified analyses. spatialMNN enables unified clustering to identify spatial domains across many samples, and is implemented with popular data structures, Seurat v5 object [34], which is used for storing SRT datasets and clustering results.

## 4 Methods

### 4.1 Summary of the spatialMNN algorithm

#### Preprocessing, dimensionality reduction, and spatial graph construction

The spatialMNN workflow begins with preprocessing steps (**Extended Data Figure 1**). There are two options for these steps. In one option, for each tissue sample, spatialMNN uses GLM-PCA [35] to perform dimensionality reduction. We use this approach because GLM-PCA has been shown to avoid distortions and potential false discoveries associated with normalization and log_2_-transformed data that is highly sparse [35]. Specifically, we use the `nullResiduals()` function from the *scry* R package [36] to obtain a set of a deviance residuals, which are used as input to principal components analysis (PCA) to calculate a set of top principal components (PCs) with a default of 30 PCs. Alternatively, a user has the option to identify highly variable genes and use the standard PCA approach to perform dimensionality reduction, if desired. Either way, the output from the dimensionality reduction step is a new matrix Y^*M*×*N*^ containing *M* spots/cells and *N* PCs.

Next, using the euclidean distance of the spatial coordinates of spots/cells, the spatialMNN algorithm constructs a *k*-Nearest Neighbors (kNN) graph (referred to as a ‘spatial graph’) using the `kNN()` function from the *dbscan* R package [37, 38]. The parameter *k* is flexible (default *k* = 6) can be decided from SRT platforms or cell density (**Extended Data Figure 2**). We denote the kNN graph as *G* =*< V, E >*, where *V* represents the set of all spots/cells and *E* the set of edges connecting each node *u* ∈ *V* with its *k* nearest neighbors. The edge weights are calculated based on Pearson’s correlation scores between the PCs of spot pairs (default), which works well for most types of data sets. Alternatively, the number of Shared Nearest Neighbors (SNN) in the PC space is useful for more sparse data sets, such as cases where more genes have zero expression (zero-inflated), or where the weight calculations based on the Pearson’s coefficient may fail.

#### Smoothed edge pruning

Next, spatialMNN prunes (or removes) edges either (i) using a fixed threshold (removing edges below 0.6 as a default) or (ii) alternatively users can use a smoothed edge pruning algorithm. A graphical summary of the algorithm is provided (**Extended Data Figure 2**) and detailed below, but the motivation of the smoothed version of edge pruning is to minimize the effects of noisy expression at the spot/cell level by considering adjacent edges of a specific edge *e* when calculating the *e*^*th*^ edge weight.

Our smoothed edge pruning algorithm aims to identify adjacent point sets for each edge by assigning edge weights and determining whether an edge should be pruned, while also identifying biologically significant boundaries. Specifically, for a given edge *e*, the connecting nodes (*u, v*), and the set of (*u, v*)’s *q* nearest neighbors in spatial domain denoted as *V*_*u*_ and *V*_*v*_, the edge weight is computed as follows:

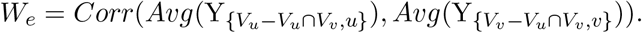

where Y^*M*×*N*^ is a matrix containing *M* spots/cells and *N* PCs.

Next, the edges with weight below the user defined threshold (default 0.6, can be determined from the distribution of edge weights) (**Extended Data Figure 3**) are pruned to form the final graph for partitioning in the next step.

#### Partitioning the spatial graph as a form of data reduction

Following edge pruning, the spatial graph undergoes partitioning via the Louvain algorithm [39] set to a high resolution (∼ 10), resulting in 100-200 small niches for each tissue sample. These niches act as a set of anchor points for the sample, but also shrinks the data from potentially thousands of spots/cells to hundreds of small niches. Niches with too few spots/cells (default *n<* 5) will be merged to nearby niches that most of the neighbors belong to. Using these niches, we further reduce the raw gene expression counts matrix by pseudobulking counts across all spots/cell within a given niche/cluster.

#### Mutual nearest neighbors on the spatial niches

Next, we construct a mutual nearest neighbors [8] graph across the niches identified within each tissue sample. Using the pseudobulked (across spots within a niche) gene expression data, the spatialMNN algorithm will first find each niche’s *k*-nearest neighbors in every sample except itself (or include itself if the user wants to detect sample-specific cell types). Suppose there are a pair of niches from different samples, *u* from sample 1 and *v* from sample 2. The algorithm will find the set, *V*_*u*_, which includes *u*’s k-nearest neighbors in sample 2 in PCA space and, *V*_*v*_ in the same way. Only if *u* ∈ *V*_*v*_ and *v* ∈ *V*_*u*_, will this pair of niches be connected in the MNN graph.

In general, the above approach applies to cells or tissue types that are shared by at least two samples, but if there is a sample-specific cell type, these nodes will not be added to the MNN network and cannot be correctly grouped together, as there are no mutual nearest neighbors in other samples. Therefore, it is possible to consider finding mutual neighbors within the sample, so that the niches contain unique cell type of the sample that can be successfully identified and grouped. After constructing the MNN graph, the graph will also be partitioned using Louvain clustering, but with a lower resolution (∼ 1). Finally, the clustering label will be assigned to all spots within a labeled niche.

#### Parallelizing the spatialMNN algorithm

The parallel computing capability of spatialMNN leverages the R *parallel* package to enhance performance. Due to spatialMNN’s two-stage design (**Extended Data Figure 1**), parallelization is straightforward. In the first stage, since the pre-processing and spatial graph construction steps for each sample are independent, different samples can be assigned to separate cores for processing. Each sample will be copied only once. For datasets with large sample sizes, this approach significantly reduces processing time while keeping memory consumption at a minimal level. In the most extreme case, where the number of available cores matches or exceeds the number of samples, using the parallel version of spatialMNN requires only double the memory for the first stage. To enable parallelization, simply set the `core_num` parameter to an integer greater than one in the `stage_1` function.

#### Resolve batch effects in large samples

In the second stage of spatialMNN, batch effects are addressed during dimensionality reduction and clustering steps. By using the Pearson residuals approximation to GLM-PCA, spatialMNN partially mitigates batch effects in the data [35]. Additionally, the use of mutual nearest neighbors (MNN) in the clustering process further reduces the impact of batch effects. By default, the MNN construction process only connects similar nodes between different samples, preventing over-connection of nodes within the same sample [8], which could otherwise result from batch effects. However, spatialMNN includes an option that allows users to enable connections within the same sample if necessary. This option is particularly useful when dealing with small sample sizes or when certain samples contain unique cell or tissue types, thereby improving clustering quality.

### 4.2 Datasets

SpatialMNN can be applied to either (i) image-based, targeted, in situ transcriptomic profiling at a molecular and single-cell resolution or (ii) non-targeted RNA capture and sequencing approaches [4].

#### 4.2.1 Mouse brain STARmap data

Gene expression counts were downloaded from Wang et al. [25] generated on the STARmap (spatially resolved transcript amplicon readout mapping) platform profiling the prelimbic area in mouse brain across *N*=3 tissue sections, 166 genes, and *n*=3,190 cells. Cell labels were assigned *a priori* to each cell in the original publication. We note that the BayesSpace algorithm could not be run on this dataset because it is not applicable to image-based *in situ* technologies profiling at a molecular and single-cell resolution.

#### 4.2.2 Visium human dorsolateral prefrontal cortex data

Gene expression counts were downloaded from Globus as described in [26]. These data were generated on the 10x Genomics Visium Spatial Gene Expression platform profiling postmortem dorsolateral prefrontal cortex (DLPFC) in human brain across *N*=12 tissue sections, 33,538 genes, and *n*=47,681 spots. Spatial domains were manually annotated at the spot-level *a priori* in the original publication.

#### 4.2.3 MERFISH mouse frontal cortex and striatum data

Gene expression counts were downloaded from [CZI CELLxGENE Collections portal](https://cellxgene.cziscience.com/collections/31937775-0602-4e52-a799-b6acdd2bac2e), which were published in Joung et al. [27]. These data were generated on the MERFISH platform profiling frontal cortex and striatum in mouse brain across *N*=31 tissue sections, 374 genes, and *n*=378,918 cells. Cell labels were assigned *a priori* to each cell in the original publication. We note that the BayesSpace algorithm could not be run on this dataset because it is not applicable to image-based *in situ* technologies profiling at a molecular and single-cell resolution. Further, BASS could not be run on this dataset due to out-of-memory issue.

#### 4.2.4 STARmap PLUS Alzheimer’s Disease mouse brain data

Gene expression counts were downloaded from Zeng et al. [28] generated on the STARmap PLUS (protein localization and unlimited sequencing) platform profiling cortical and hippocampal regions in TauPS2APP triple transgenic mice, an established AD mouse model that exhibits both amyloid plaque and tau pathologies. Gene expression and protein levels were measured in *N*=4 tissue sections from 8- and 13-month-old TauPS2APP mouse and normal mouse brains in the context of extracellular amyloid-*β* (A*β*) plaques and intracellular hyperphosphorylated tau accumulation. A targeted list of 2,766 genes were profiled measuring *n*=35,094 cells and 2 proteins.

#### 4.2.5 MERFISH mouse hypothalamic preoptic region data

Gene expression counts were downloaded from Moffitt et al. [33] generated on the MERFISH platform profiling mouse hypothalamic preoptic regions from 6 social behaviour types in adult C57Bl6/J 16 female and 20 male mice. A targeted panel of 161 genes was measured in *n*=1,027,848 cells from *N*=181 tissue sections, and cell labels were assigned *a priori* to each cell in the original publication. After filtering out cells with “Ambiguous” label, 874,768 cells were used in the benchmark.

#### 4.2.6 Visium human hippocampus data

Gene expression counts were downloaded from Nelson et al. [32] generated on the 10x Genomics Visium Spatial Gene Expression platform postmortem hippocampus (HPC) in human brain across *N*=31 tissue sections from 10 neurotypical donors (2-5 Visium slides per donor), 31,483 genes, and *n*=150,917 spots.

#### 4.2.7 Generation, preprocessing, and analysis of simulated data

As the data set that is manually annotated still may have an inaccurate or false annotations, we also considered simulating data to assess the accuracy of each method, because we can define the ground truth for each sample. Further, since we can control the size of the generated sample we can evaluate the efficiency of the algorithm, including the amount of time and memory that is consumed during the operation. Time and memory consumption are both of utmost importance as they determine how well an algorithm can scale when facing the challenge of larger datasets in the future.

The data generation involves two phases (**Figure 5**). The first phase determines the spot array (the position for each spot), the tissues and cell types contained in the generated data and their spatial distribution. Here, a tissue is defined as a region containing multiple cell types, aligning more closely with real-world scenarios. The spatial distribution patterns of the tissues typically include layered or ringed structures, with the specific size of the distribution area determined manually. A sample usually comprises thousands of units (each unit consisting of coordinates and transcriptomic information). Each tissue can contain multiple cell types. The proportions of these cell types can be manually set and represented by constructing a probability matrix of tissue-cell type, as shown in Table 1, below.

**Table 1:**
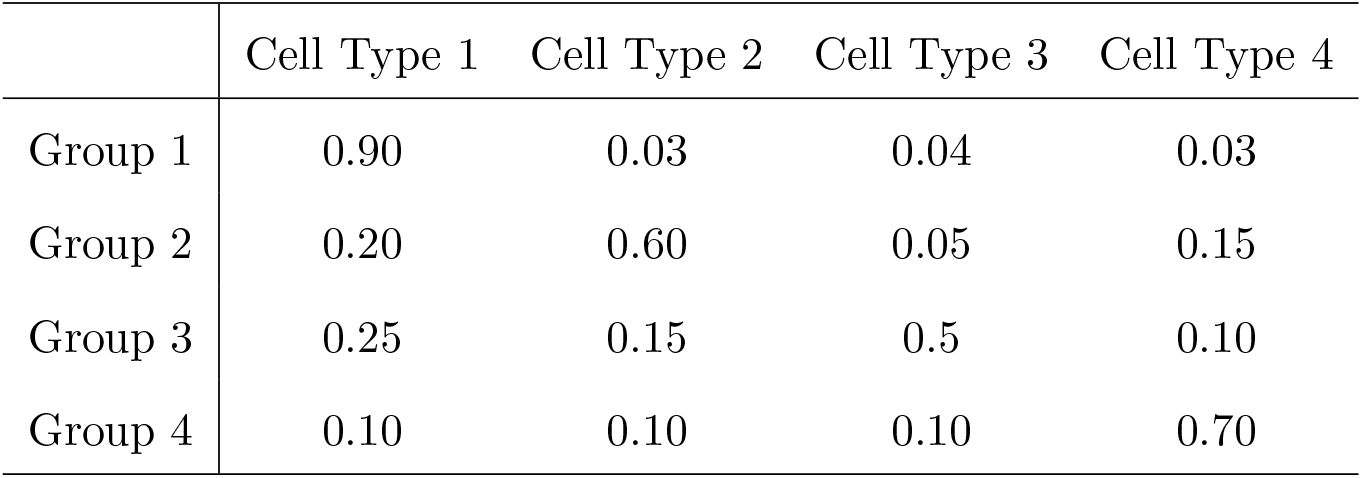
Cell Type Composition Matrix.

The second phase involves generating SRT data based on the tissue-cell type probability matrix, that is, determining the cell type or composition for the units generated in the previous phase. To reflect the differences in data generated by different technologies, in a cell-based dataset, a unit contains only one cell. In contrast, in a spot-based dataset, a unit consists of several cells, represented by averaged expression profile data. The cell types of these cells are randomly generated based on the aforementioned probability matrix. For transcriptomic data, each cell type has 200 highly expressed genes (referred to as marker genes). There are also 200 noisy genes to simulate real-world scenarios. Regarding gene expression, the marker genes of a particular cell type will be highly expressed in the corresponding cells, while all other genes maintain a zero expression level. After generation, all expression values will have an added layer of Gaussian noise to simulate real-world situations. The formula for data generation process is described below:

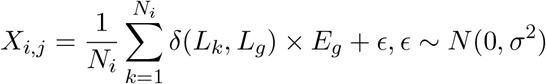

Where *i, j* stands for the index of unit and gene(feature), *N*_*i*_ is the number of cells in the unit, *L*_*k*_, *L*_*g*_ are the cell type label of cell *k* and the cell type which *g*’s marker gene list belongs to, and *δ* is the Kronecker delta function which outputs 1 when the two inputs are equal and 0 otherwise.

### 4.3 Existing computational methods

We compared spatialMNN to existing methods that integrate multiple SRT tissue samples and perform unsupervised clustering. In our benchmark evaluations, we used default parameters obtained from the developer-provided tutorials or publications, unless noted below.

#### 4.3.1 BASS

BASS [13] performs unsupervised clustering across multiple SRT samples either at a single-cell resolution or for spatial domain detection using a Bayesian hierarchical modeling framework. It takes as input a list of raw gene expression matrices and spatial coordinates of cells or spots, along with sample information like the expected number of cell types (C) and spatial domains (R). These parameters are not optional and the user needs to provide some values for these when creating BASS object. BASS provides the outputs including the inferred cell type cluster labels, spatial domain labels, and the cell type proportions inside each spatial domain. In the benchmark, we used the spatial domain labels (*z*) to calculate the accuracy metrics. We used the recommended parameters in benchmark: For STARmap datasets, the region number is set to 4 and top 10 PCs are used in the preprocessing step. Region number and the number of PCs are set to 7 and 20 respectively for DLPFC dataset. The software is available on GitHub (github.com/zhengli09/BASS) and we used version 1.6.5.

#### 4.3.2 PRECAST

PRECAST [14] is a probabilistic framework designed for joint analysis of multiple spatial transcriptomics datasets. It integrates spatial coordinates and gene expression profiles to identify spatial domains while accounting for batch effects across samples. PRECAST takes as input the spatial locations and gene expression matrices of multiple SRT datasets, optionally incorporating covariates or batch labels. As output, it provides spatially coherent domain clusters across datasets while simultaneously correcting for technical variations, improving the biological interpretability of spatial domain assignments. The number of cluters is set to 4 (STARmap), 7 (DLPFC), and 8 (MERFISH) respectively. The software is available on CRAN (cloud.r-project.org/web/packages/PRECAST/index.html) and we used version 1.1.0.016.

#### 4.3.3 BayesSpace

BayesSpace [11] enhances spatial resolution in SRT data by refining spatial domain boundaries using a Bayesian modeling approach. It models spatial expression data through a mixture model that accounts for neighboring relationships between spots to improve the detection of fine-grained spatial domains. BayesSpace takes as input spatial gene expression data from a standard SRT experiment and optional neighborhood information. As output, it provides high-resolution spatial domain assignments, offering a refined clustering of the spatial transcriptomics data that accounts for local dependencies and spatial structure. Based on its recommended type of SRT platforms, we only run BayesSpace on the DLPFC and simulated datasets. The first 50 PCs are used for spatial preprocessing and the number of clusters is set to 7 and 4 respectively. The software is available on Bioconductor (www.bioconductor.org/packages/release/bioc/html/BayesSpace.html), and we used BayesSpace version 1.6.0.

#### 4.3.4 BANKSY

BANKSY [15] performs unsupervised clustering at spatial domain or cell type level using the gene expressions of cells as well as their neighbourhood representations built using weighted average of neighbourhood gene expressions and an azimuthal Gabor filter. It takes a SpatialExperiment object [40] as input, and the cluster labels generated by the algorithm are added to the metadata in the object. We followed BANKSY preprocessing steps such as trimming the datasets to save memory. For datasets with samples from different sources, additional preprocessing steps such as staggering spatial coordinates to avoid overlap between locations from different samples and running harmony for batch correction were performed. Whole transcriptome spatial transcriptomics datasets, like the DLPFC dataset, were subjected to feature selection using FindVariableFeatures function (nfeatures = 2000) of Seurat [41], prior to batch correction and in accordance with BANKSY protocol. The selection of BANKSY parameters such as neighbourhood size (k geom), mixing parameter (lambda), Leiden clustering resolution (res), and number of PCA dimensions (npcs) depended on the type of input dataset. We used BANKSY recommended values for domain-level clustering – for DLPFC dataset k geom = 18, lambda = 0.2, res = 0.55, npcs = 20; for STARmap dataset k geom = 30, lambda = 0.8, res = 0.8, npcs = 50; and for MERFISH dataset k geom = 30, lambda = 0.8, res = 1, npcs = 20. The software is available on GitHub (github.com/prabhakarlab/Banksy), and we used BANKSY version 1.0.0.

#### 4.3.5 SLAT

SLAT [16] is a graph convolutional network-based alignment tool capable of mapping multiple samples from different technologies and modalities using a graph adversarial matching algorithm. For multisample alignment, SLAT requires a list of AnnData objects [42] arranged in the correct order for 3D reconstruction of original tissue. The output is a list of best matching cell indices from each pair of samples aligned. For benchmarking purpose, we added a Leiden clustering step prior to SLAT alignment, such that clustering was performed on the first sample in the AnnData object list, followed by multisample alignment, and then sequential borrowing of cluster labels from first to second sample, second to third sample, and so on. To ensure fair treatment, we performed this clustering + SLAT approach using each sample in the dataset as the first clustered sample. To maximize the number of alignments in SLAT, we used a low cosine similarity cutoff of 0.3. SLAT software is available on GitHub (github.com/gao-lab/SLAT), and we used SLAT version 0.2.1.

#### 4.3.6 MENDER

MENDER [17] is a multisample, clustering method that performs spatial domain detection via a multi-range neighbourhood representation approach. MENDER takes AnnData objects as input, and it requires cell state information in the form of prior cell annotations or cluster labels to generate spatial context for each cell over a range of neighbourhood sizes. MENDER output contains an AnnData object with MENDER cluster labels. For benchmarking, we used Harmony algorithm [7] on samples from different sources to counter batch effects and Leiden clustering to generate cell state information, as recommended. MENDER parameters such as number of neighbourhood sizes (n_scales), neighbourhood mode based on the spatial technology of input dataset (nn_mode), neighbourhood size (nn_para), and final Leiden clustering resolution (target_k) were selected based on the type of input data. We followed MENDER tutorial for making these parameter selections for each dataset – for DLPFC dataset n_scales = 6, nn_mode = ‘ring’, nn_para = 6, and target_k = -0.2; for STARmap and MERFISH datasets n_scales = 6, nn_mode = ‘radius’, target_k = -0.5/8, and nn_para = 150/15 (STARmap/MERFISH). The software is available on GitHub (github.com/yuanzhiyuan/MENDER), and we used MENDER version 1.1.

#### 4.3.7 Louvain method for community detection

Louvain clustering [39], which is embedded in Seurat v3 [41], is a graph-based approach for detecting clusters in single-cell data. It operates by constructing a shared Nearest-neighbor (SNN) graph and optimizing the modularity to detect communities or clusters. Louvain takes as input a precomputed KNN graph derived from an expression matrix, along with a resolution parameter that controls cluster granularity. The output is a set of clusters representing either cell types or spatial domains, with flexibility in adjusting the resolution to obtain finer or broader clusters. As recommended by Seurat tutorial, the raw count matrices of all datasets are used as input. Followed by scaling, normalization and HVG selection, the dimension of the expression matrices are further reduced by PCA. For SNN construction, we used top 10 PCs and default resolution (0.1) for clustering. The software is available on CRAN (cran.r-project.org/web/packages/Seurat/index.html), and we used Seurat version 5.0.3.

### 4.4 Overview of metrics used for evaluation

#### Performance metrics

We used the adjusted Rand index (ARI) to compare the overlap of the manual annotations at a spot/cell resolution to a predicted cell type or spatial domain. An ARI of 0 corresponds to random labeling and an ARI of 1 corresponds to perfect agreement. We used the `adjustedRandIndex()` function in the *mclust* R package [43] (version 6.0.0). We also used normalized mutual information (NMI) to evaluate the accuracy using our simulated SRT samples. A NMI of 0 corresponds to no mutual information and 1 corresponds to perfect correlation. We used the `NMI()` function in the *aricode* R package [44] (version 1.0.3).

#### Spatial feature related metrics

In Figure 3 (the analysis of AD mouse datasets), we used the co-localization index to demonstrate the spatial relationship of identified domains with A*β* plaque. The index of a certain domain is calculated by: 1. counting the nearby plaques (*<*20 *µm*) for each cell; 2. averaging the plaque number by domains.

#### Computational environment

In R related benchmark, we evaluated both the max or peak memory used and computing time using the `peakRAM()` function in the `peakRAM` R package [45] and the `Sys.time()` function in base R (version 4.3/4.4), respectively. We performed all benchmark evaluations on a high performance computing (HPC) cluster with 8 cores and a max of 128GB memory.

### 4.5 Software availability

spatialMNN is freely available as an R package on GitHub (github.com/Pixel-Dream/spatialMNN) and will be submitted to Bioconductor. Code to reproduce all preprocessing, analyses, and figures in this manuscript is available from GitHub at github.com/Pixel-Dream/spatialMNN_analysis. We used spatialMNN version 0.99.0 for the analyses in this manuscript.

## 5 Back Matter

## Acknowledgments

We thank the maintainers of the Joint High Performance Computing Exchange (JHPCE) compute cluster at Johns Hopkins Bloomberg School of Public Health for providing essential computing resources. We also thank Kasper Hansen, Haotian Shi (Shanghai Jiao Tong University), Zefang Tang (Broad Institute), Yifan Wu (Wuhan University), Yuqing Yang (Peking University) and members of the Hicks lab for their constructive comments and suggestions. We thank members of the Sydney Precision Data Science Centre for their engagement.

## Author Contributions Statement

HZ conceptualized the spatialMNN methodological framework with suggestions from SCH, BG, PP, and SG. The formal analysis and benchmarking was led by HZ and PP. Figures were designed by HZ. Software was developed by HZ with documentation support from CH and input from other co-authors. SG and SCH administered and supervised the project. Data visualizations were developed by HZ. HZ and SCH wrote the original draft of the text and all co-authors edited and approved the final manuscript.

