## Extended Data Figures 1-6 for "Spatial mutual nearest neighbors for spatial transcriptomics data"

### Contents

1. **Extended Data Figures 1-6**

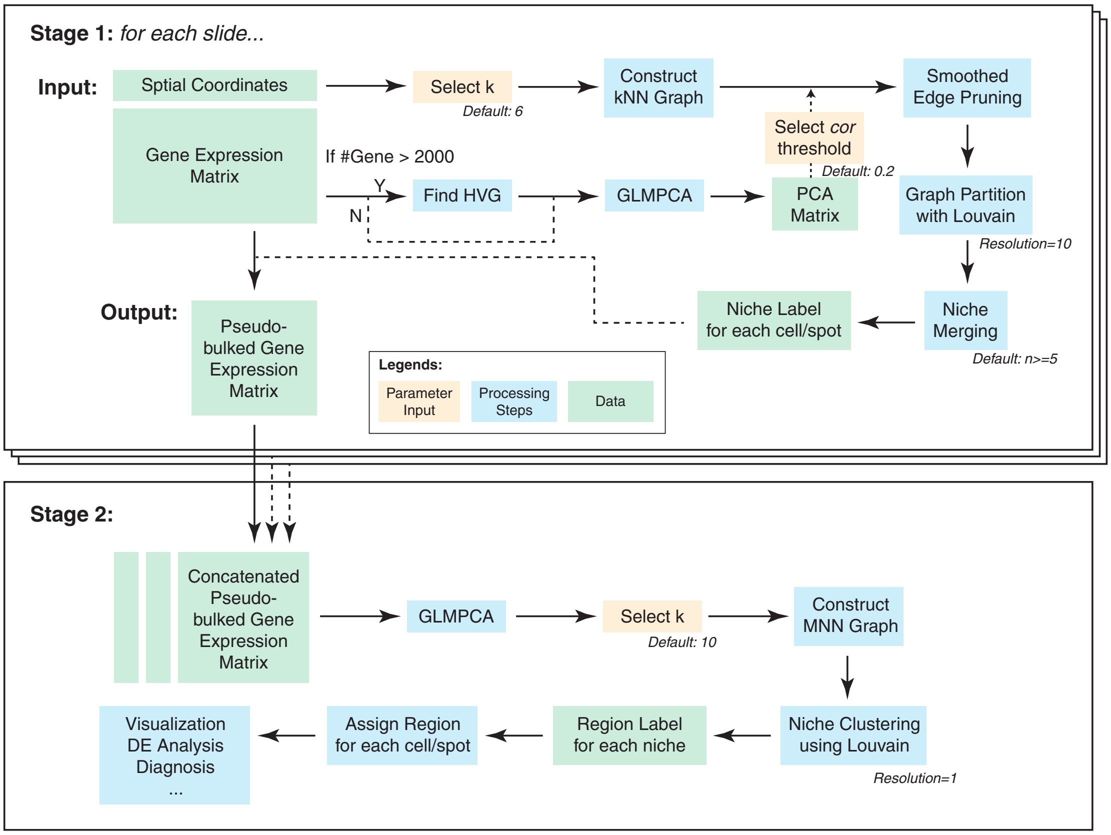


**Extended Data Figure 1: The detailed schematic of the two-stage pipeline used in spatialMNN.** Stage 1 begins with input data consisting of spatial coordinates and gene expression matrices for each slide. Initially, a k-nearest neighbor (kNN) graph is constructed (default $k = 6$) based on spatial distances. Highly variable genes (HVGs) are identified if the number of genes exceeds 2000. Generalized Linear Model Principal Component Analysis (GLMPCA) is applied to reduce dimensionality, producing a PCA matrix. A correlation threshold (default $cor = 0.2$) is set, and smoothed edge pruning is performed to refine the graph. The correlation threshold can be determined from the weight distribution of all edges. Typically the threshold will be higher if applying SpatialMNN on spot-based platforms. The graph is then partitioned using the Louvain algorithm with a resolution parameter of 10, and niche merging is conducted (default $n < 5$). The output of this stage is a pseudo-bulked gene expression matrix and niche labels for each cell or spot. Stage 2 involves the concatenation of pseudo-bulked gene expression matrices across slides. GLMPCA is applied again for dimensionality reduction, followed by kNN graph construction (default $k = 10$) and the construction of a mutual nearest neighbor (MNN) graph. Niche clustering is performed using the Louvain algorithm (resolution = 1), resulting in region labels for each niche. Downstream analysis like visualization, differential expression analysis, and diagnosis insights for each region can be done using spatialMNN output labels.

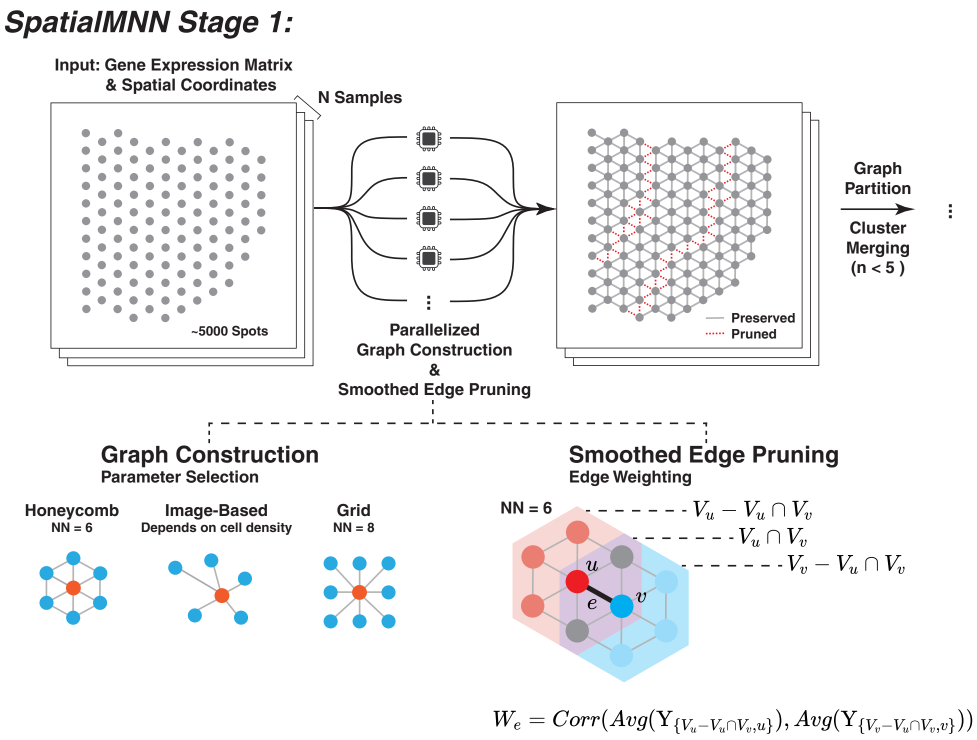


**Extended Data Figure 2: Overview of the smoothed edge pruning algorithm**. Parallelized graph construction and edge pruning are applied independently to each sample, with input of gene expression matrices and spatial coordinates. Various graph topologies, such as honeycomb (suitable for 10X Visium, NN = 6) and grid (NN = 8), are used to represent spatial relationships between spots. For image-based SRT technologies, such as MERFISH and STARmap, the NN is dependent on cell density. Smoothed edge pruning is then performed by calculating the correlation of average gene expression between neighboring spots, where $W_e$ is the correlation between the gene expression of neighboring nodes after excluding their shared neighbors. This process removes weak or noisy connections, preserving only significant spatial relationships. The refined graph is subsequently partitioned and clusters are merged (typically $n < 5$), facilitating robust identification of spatial niches.

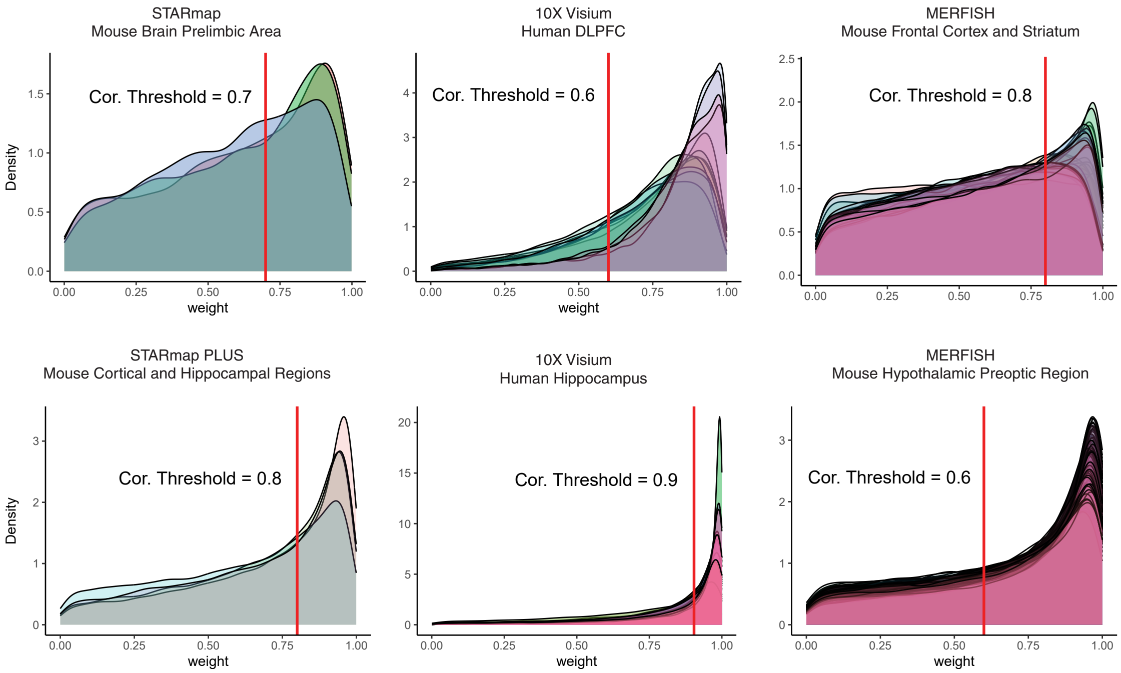


**Extended Data Figure 3: Edge Weight Distribution and threshold used in spatialMNN.** The weight distribution of the spatial coordinates-based graph constructed in the stage 1, the distribution of each sample is displayed in different colors, and the threshold value used for the graph pruning in stage 1 is marked with a red vertical line. The value is determined according to the elbow point of the overall weight distribution.

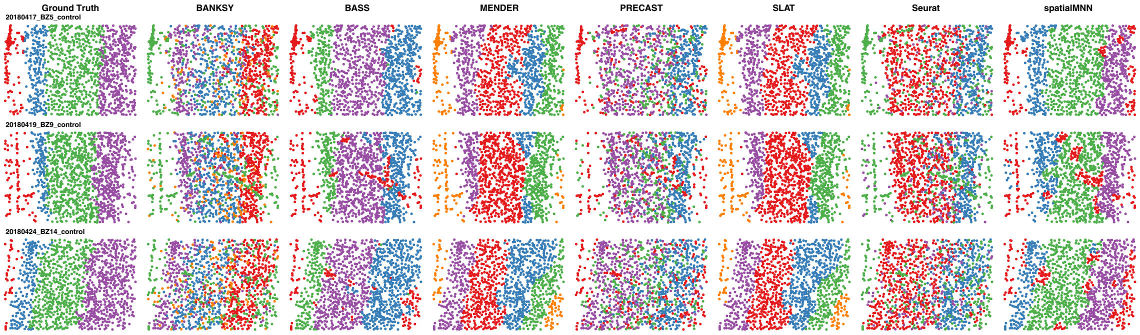


**Extended Data Figure 4: Benchmark results of STARmap mouse brain prelimbic area data.**

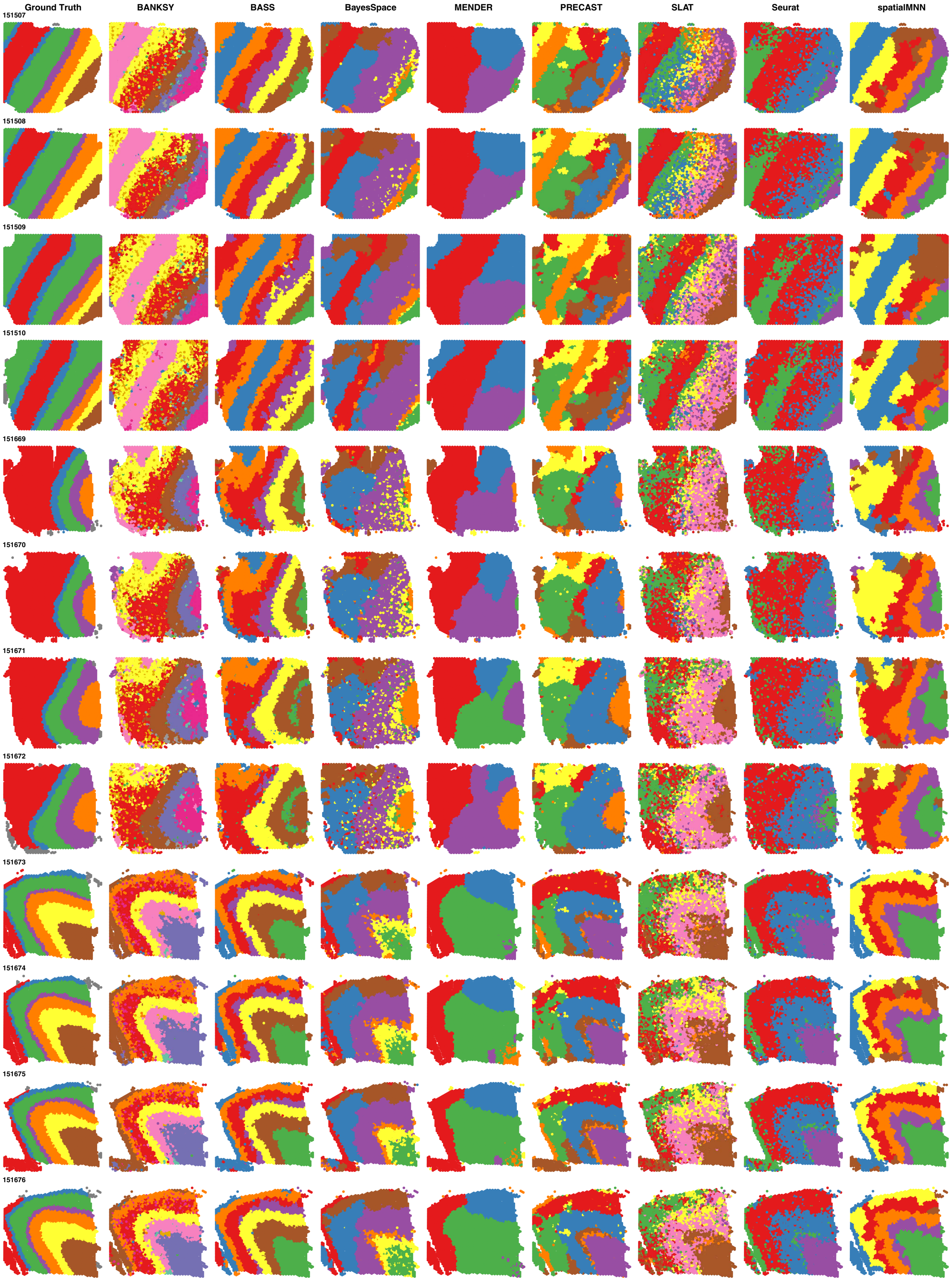


**Extended Data Figure 5: Benchmark results of Visium human dorsolateral prefrontal cortex data.**

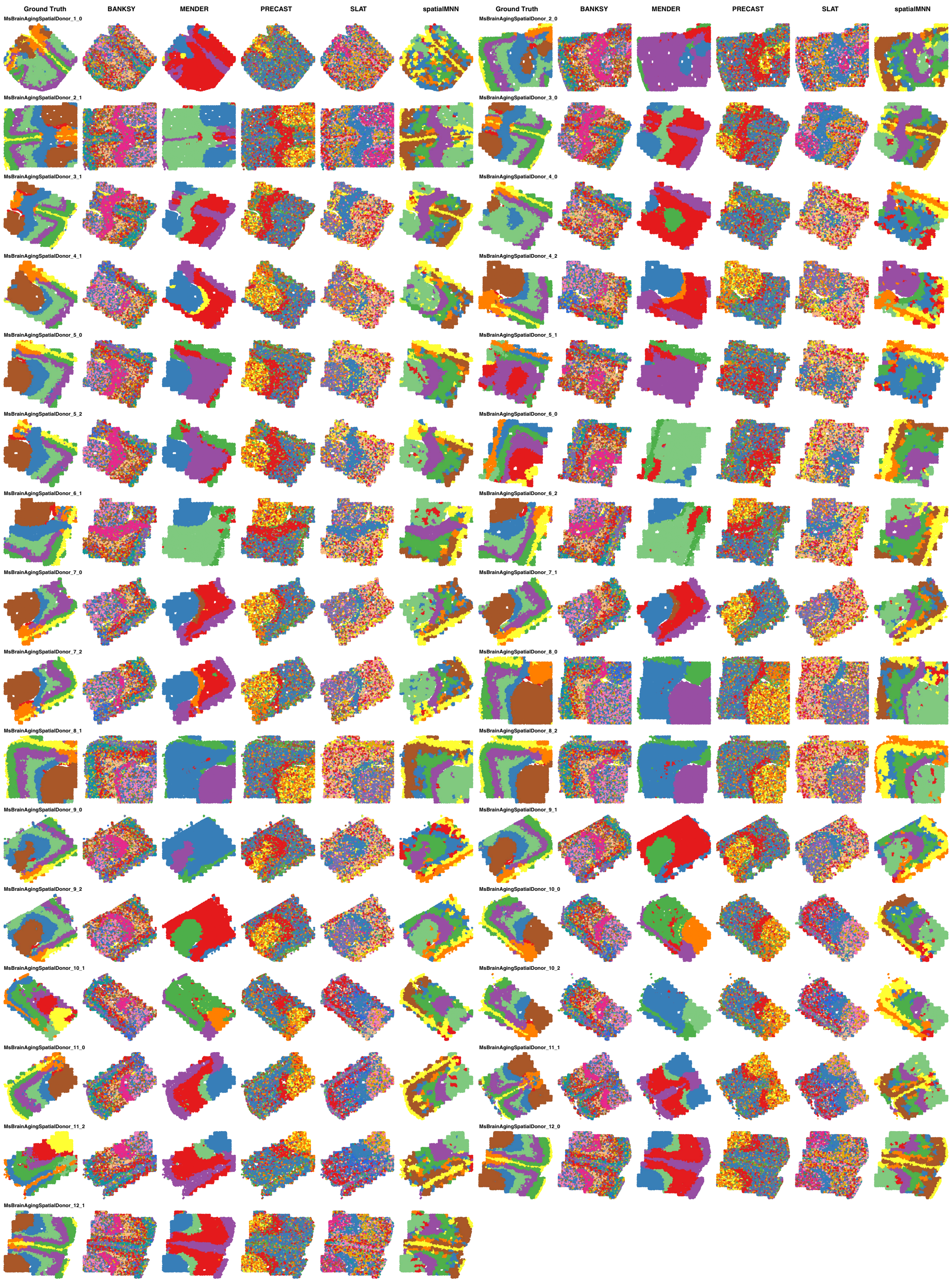


**Extended Data Figure 6: Benchmark results of MERFISH mouse frontal cortex and striatum data.**
